# Molecular Glue-Design-Evaluator (MOLDE): An Advanced Method for In-Silico Molecular Glue Design

**DOI:** 10.1101/2024.08.06.606937

**Authors:** A S Ben Geoffrey, Deepak Agrawal, Nagaraj M Kulkarni, G Manonmani

## Abstract

Protein function modulation using small molecule binding is an important therapeutic strategy for many diseases. However, many proteins remain undruggable due to lack of suitable binding pockets for small molecule binding. Proximity induced protein degradation using molecular glues has recently been identified as an important strategy to target undruggable proteins. Molecular glues were discovered serendipitously and as such currently lack an established approach for in-silico driven rationale design. In this work, we aim to establish an in-silico method for designing molecular glues. To achieve this, we leverage known molecular glue-mediated ternary complexes and derive a rationale for in-silico design of molecular glues. Establishing an in-silico rationale for molecular glue design would significantly contribute to the literature and accelerate the discovery of molecular glues for targeting previously undruggable proteins. Our work presented here and named as Molecular Glue-Designer-Evaluator (MOLDE) contributes to the growing literature of in-silico approaches to drug design in-silico literature.

## Introduction

Proteins play important functional roles in the human biological system and in disease their function becomes either up or down requiring them to be modulated in function using some therapeutic means of intervention. However, only about 20% of the human proteome is amenable to modulation of function using small molecules and the remaining proteins are considered ‘undruggable’ due to lack of suitable binding pockets for small molecules [1]. One promising solution is the modulation of Protein-Protein Interactions (PPIs) using small molecules. PPIs are critical for cellular processes but targeting them with traditional small molecules has proven difficult [2]. Recently, a new class of compounds called molecular glues has emerged. These small molecules can enhance PPIs by bringing proteins into proximity, thereby stabilizing and modulating their interactions [3]. However, designing molecular glue remains a challenge and there are no established in-silico methods in the literature for rational molecular glue design [3–6]. In this research we address this gap by using well-established as well as novel in-silico methods for small molecules including New Chemical Entity (NCE) generation, NCE optimization, molecular docking, and advanced molecular dynamic simulations for design of molecular glues.

While in-silico approaches for molecular glue design have not been reported in literature, in-silico approaches to design hetero-bifunctional molecules such as PROTACs to induce PPI have been reported [7–11]. The hetero-bifunctional molecule in a PROTAC must have two functional domains that interact with the two different proteins and bring them together to induce the PPI. In contrast, molecular glues are a small molecule without a linker which directly “glue” with the two target proteins. Molecular glues offer a few important advantages over PROTACs. Mainly because molecular glues are small molecule like, they do not suffer from the bioavailability and cell permeability issues associated with the larger molecules like PROTACs. Consequently, it is more desirable to use molecular glues to induce PPI especially in applications such as protein degradation. Hence it is important to have a rational in-silico design approach for molecular glues which can significantly help with designing effective molecular glues with the help of in-silico methods. In our previous work, we have illustrated an in-silico rationale for the design of PROTACs [11].

Drawing from that experience, we present an in-silico approach for de novo molecular glue design. This research aims to bridge the gap and provide rational methods for designing effective molecular glues.

The in-silico design rational was developed by studying molecular glue mediated ternary complexes in RCSB Protein Data Bank (https://www.rcsb.org/) and a retrospective validation of the developed approach was carried out to reproduce experimentally known aspects of molecular glue mediated ternary complexes. This validation demonstrates the validity of the in-silico approach. The aspects involved in the retrospective validation of the approach are as follows:

1. Reproducing molecular glue binding mode and structure of ternary complex formed therein.
2. Reproducing the thermodynamic favorability of molecular glue mediated ternary complexation through theoretical ΔG calculations.
3. Differentiating the stability of the molecular glue mediated ternary complex.

In their review, Huan et. al [12], provide a list of PDB IDs associated with known molecular glue mediated ternary complexes. In our work, we have used the molecular interactions involved in all the PDB IDs reported by Huan et.al. to understand the types of interactions involved in molecular glue mediated ternary complexes. Additionally, we have identified systems to develop various aspects of our validation and design method. The PDB ID: 6TD3 served as the system for developing our methodology to identify molecular glue binding poses and determine the structure of ternary complex formed. Systems associated with PDB IDs: 8G46, 7TE8, 6TD3 were used for theoretical ΔG calculations, providing a rational for the thermodynamic favorability of molecular glue mediated ternary complexation. Further, systems associated with PDB IDs: 6H0G and 7BQV were employed to develop an in-silico methodology for differentiating the stability associated with molecular glue mediated ternary complex. Lastly, using the developed methodology we designed a new molecular glue for the system represented by PDB ID: 6TD3.

## Methodology

### 1. The molecular glue binding mode and structure of ternary complex formed therein

Methodological aspects that were followed to reproduce the molecular glue binding mode and structure of the molecular glue mediated ternary complex formed therein are reported here. In the chosen system with PDB ID: 6TD3, the molecular glue associated with ID RC8 recruits DNA damage-binding (DDB) protein 1 to tag and degrade Cyclin dependent Kinase 12 (CDK12). A protein-protein docking was carried out with the two proteins DDB and CDK and low energy poses with a pocket formed at the interface was identified and molecular docking of the molecular glue RC8 was carried out at the pocket formed at the interface of these two proteins DDB and CDK12. Results obtained are discussed in the ‘Results and Discussion’ section.

### 2. Molecular glue mediated favorability of the two proteins to form a ternary complex

The diversity of interactions in the list of PDB IDs associated with molecular glue mediated ternary complexes collected by Haun et. al [12] was studied using LigPlot+ [13]. Further to understand the role of the molecular glue inducing the thermodynamic favorability of the Protein-Protein interaction, bio-molecular simulations were carried out using GROMACS [14] molecular dynamics package; and ΔG calculations using gmx_mmpbsa [15] tool were carried out to understand the molecular glue induced thermodynamic favorability for the protein-protein interaction. The protein residues were parameterized using the ‘AMBER99SB-ILDN’ forcefield and the ligand was parameterized using acpype [16]. The system was solvated in cubic solvent box and the TIP3P solvent model was used, and ions were added to physiological pH. To mimic physiological temperature and pressure, the system was heated to 300K temperature and 1 BAR pressure in 100 ps NVT and NPT runs with the temperature and pressure controlled using the Berendsen thermostat/barostat. A production run of 50 nanoseconds was executed and the stability of the molecular glue mediated ternary complex was accessed through the RMSD stabilization. The stable portion of the trajectory was used for MMPBSA based theoretical ΔG calculations to understand molecular glue mediated thermodynamic favorability for ternary complexation using the gmx_mmpbsa tool [15].

Furthermore, molecular glues are known to induce protein-protein interaction favorability by inducing conformation change [12] and enhancing the protein-protein interaction. To evaluate the role of molecular glue induced conformation change and enhanced protein-protein interaction achieved therein, a wide range of conformation ensembles of protein-protein poses were generated using AlphaFlow [17] and the resulting conformation change induced enhancement of protein-protein interaction between the Apo and Holo form was analyzed using interaction energies from the SURFACE tool [18].

### 3. In-silico differentiation of the stability and ternary binding constants (K_d_) associated with the molecular glue mediated ternary complex

Free energy perturbation (FEP) calculations are routinely used in small molecule drug design to obtain reliable estimates of the absolute binding free energies (ΔG) of the small molecule binding to protein targets. Herein we use FEP binding free energy calculations to differentiate nanomolar and micromolar ternary binding constants (K_d_) associated with a molecular glue mediated ternary complex. Free energy perturbation (FEP) protocol was carried out as per earlier established protocols [19, 20], wherein the ligand is decoupled from the protein in multiple lambda steps where the coulomb and Van Der Waals interactions are turned off in a step wise manner involving several Lambda steps and the ΔG difference between the coupled and the decoupled state is then used as an estimate of the ΔG of binding. The Free Energy Perturbation (FEP) protocol was implemented in GROMACS [14] biomolecular simulation package.

Thermal Titration Molecular Dynamics (TTMD) originally developed within the context of differentiating tight and weak small binders of protein targets [24,25] was adapted for the molecular glue problem to differentiate the stability of molecular glue mediated ternary complex. A classical molecular dynamics simulation was carried out for 5 nanoseconds in an increasing temperature ramp and the Tanimoto similarity is computed between interaction fingerprint vector of the first frame and the subsequent frames. The similarity index measures the retention of interaction which is expected to be more for a strong binder as compared to a weak binder.

And lastly, Quantum Mechanical/ Molecular Mechanics (QM/ MM) calculations were carried out to estimate the QM level interaction energy between the molecular glue and the pocket residues, and thereby develop QM/MM based interaction energy as a score to differentiate ternary binding constants (K_d_) of different magnitudes associated with a strongly binding and weakly binding molecular glue mediated ternary complex. The QM/MM methodology is adapted from Wang et. al. [26] where the QM based interaction energy for a small molecule bound to a protein pocket is given by

QM based interaction energy = QM energy of complex – QM energy of ligand (molecular glue) – QM energy of pocket residues of protein

The use of QM/ MM methods serves to provide extra validation to the molecular glue design, similar to the MMPBSA and TTMD methods.

The in-silico molecular glue design method (MOLDE which is an acronym for Molecular Glue Design & Evaluator) is summarized as a flow diagram in Figure 1.

**Figure 1.**
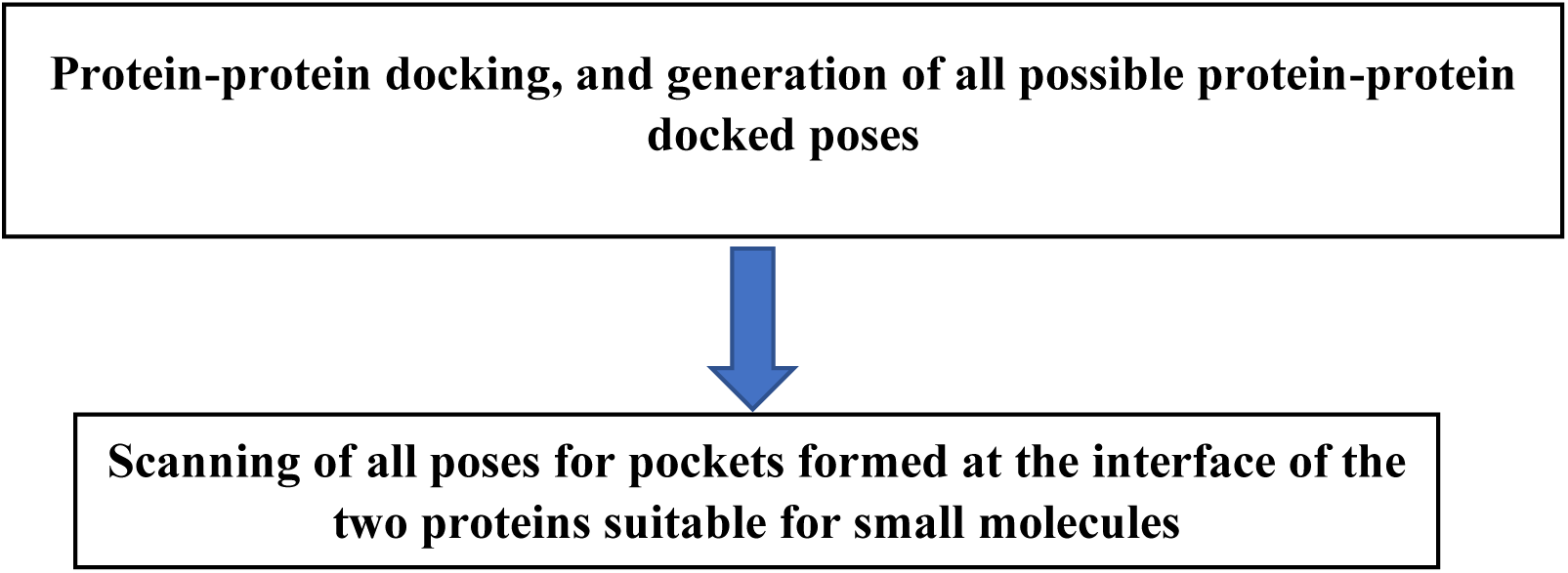
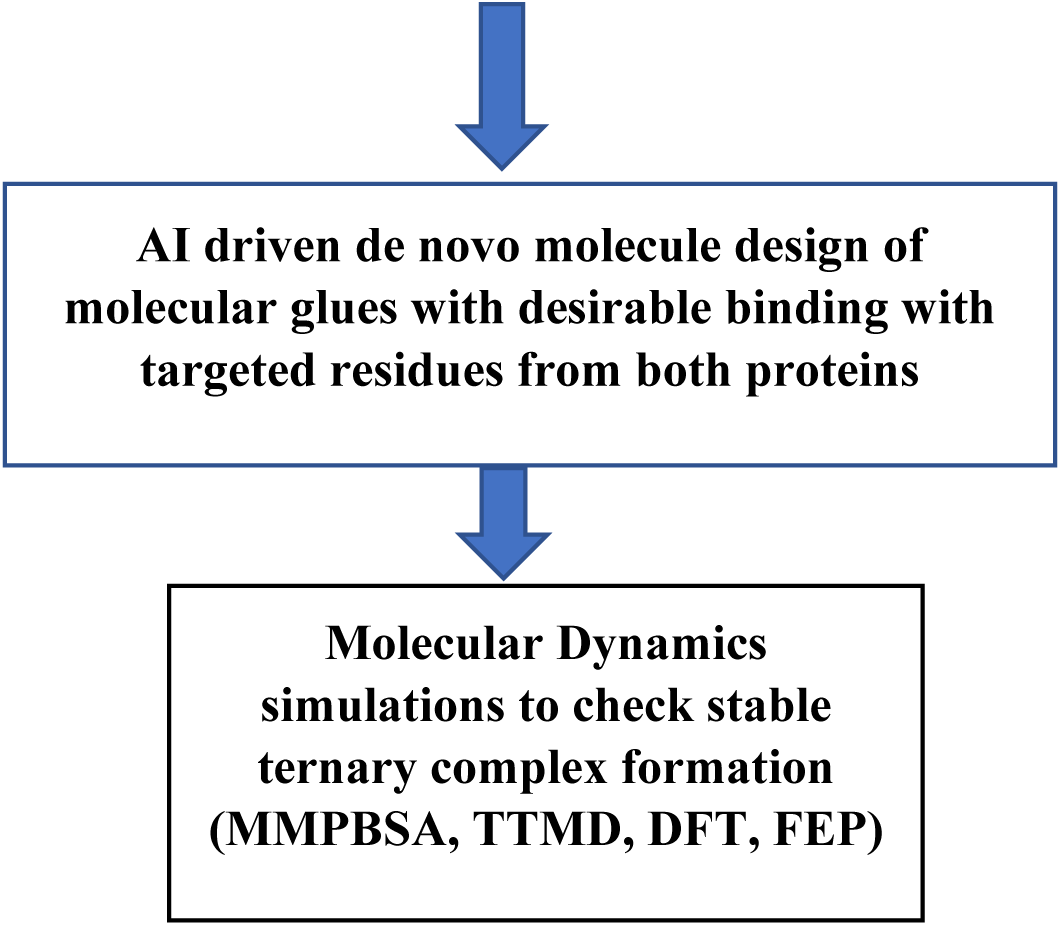
In-silico driven molecular glue design method (MOLDE: Molecular Glue Design & Evaluator)

### 4. AI driven de novo molecular glue design demonstration

Having thus developed the methodology and demonstrated its ability to reproduce the known aspects of experimentally reported molecular glue mediated ternary complexes in RCSB PDB, we apply the validated methodology to design a novel molecular glue using Generative AI for the PDB ID: 6TD3 system and results obtained are discussed below.

## Results and Discussion

### 1. Molecular glue binding mode and structure of ternary complex formed therein

In the system (PDB ID: 6TD3), the molecular glue RC8 mediates the ternary complex between the two proteins DDB1 and CDK12. Protein-Protein docking was carried out using MegaDock for the protein DDB1 and CDK12 and among the low energy poses, a pose with pocket formed at the interface that a molecular glue molecule can bind was identified as shown below and among the low energy poses as tabulated in Table 1, the lowest energy pose with pocket comparability at the interface was shortlisted for the next phase and is shown graphically in Figure 2 and Figure 3 below.

**Figure 2.**
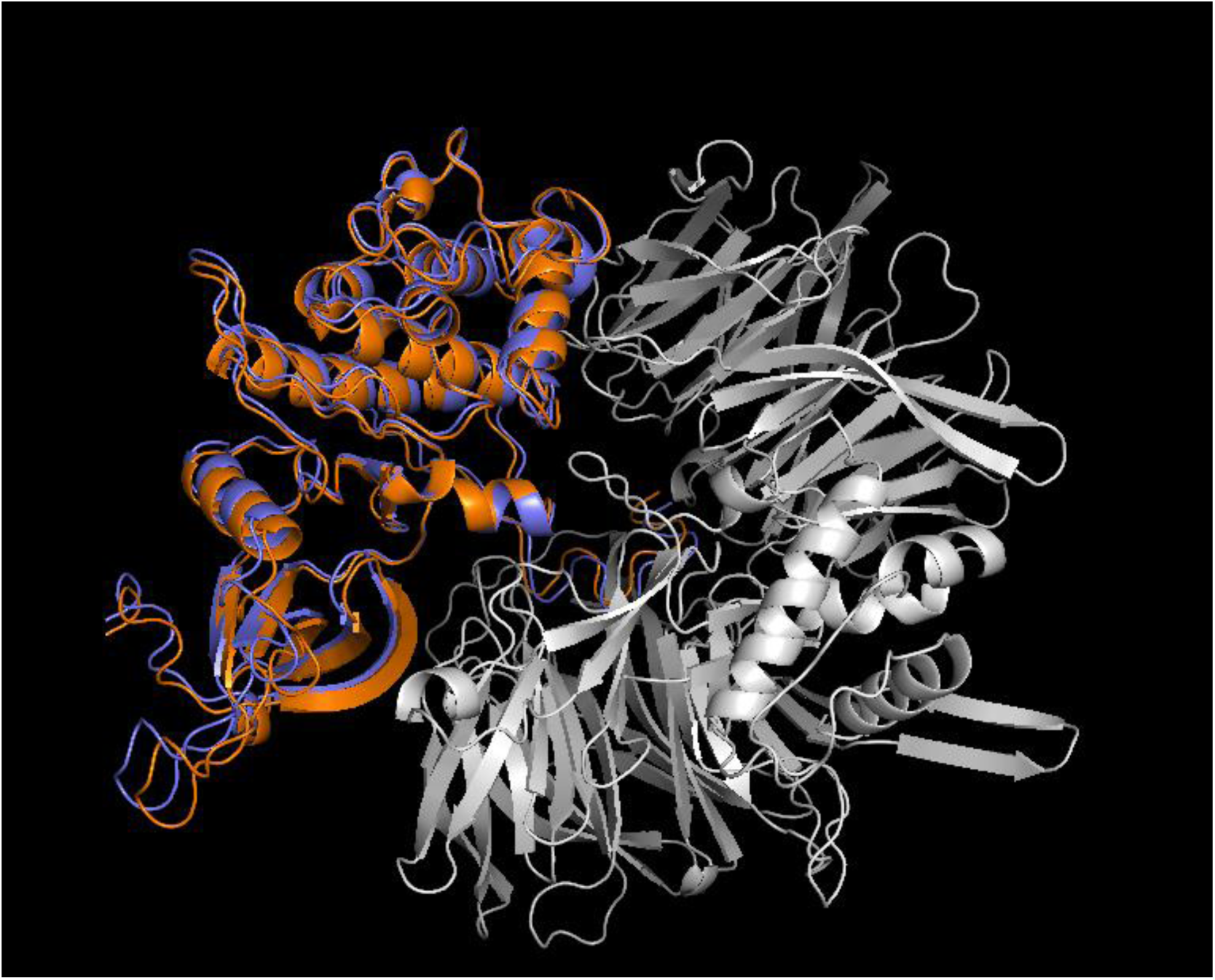
Experimental pose reproduced in protein-protein docking.

**Figure 3.**
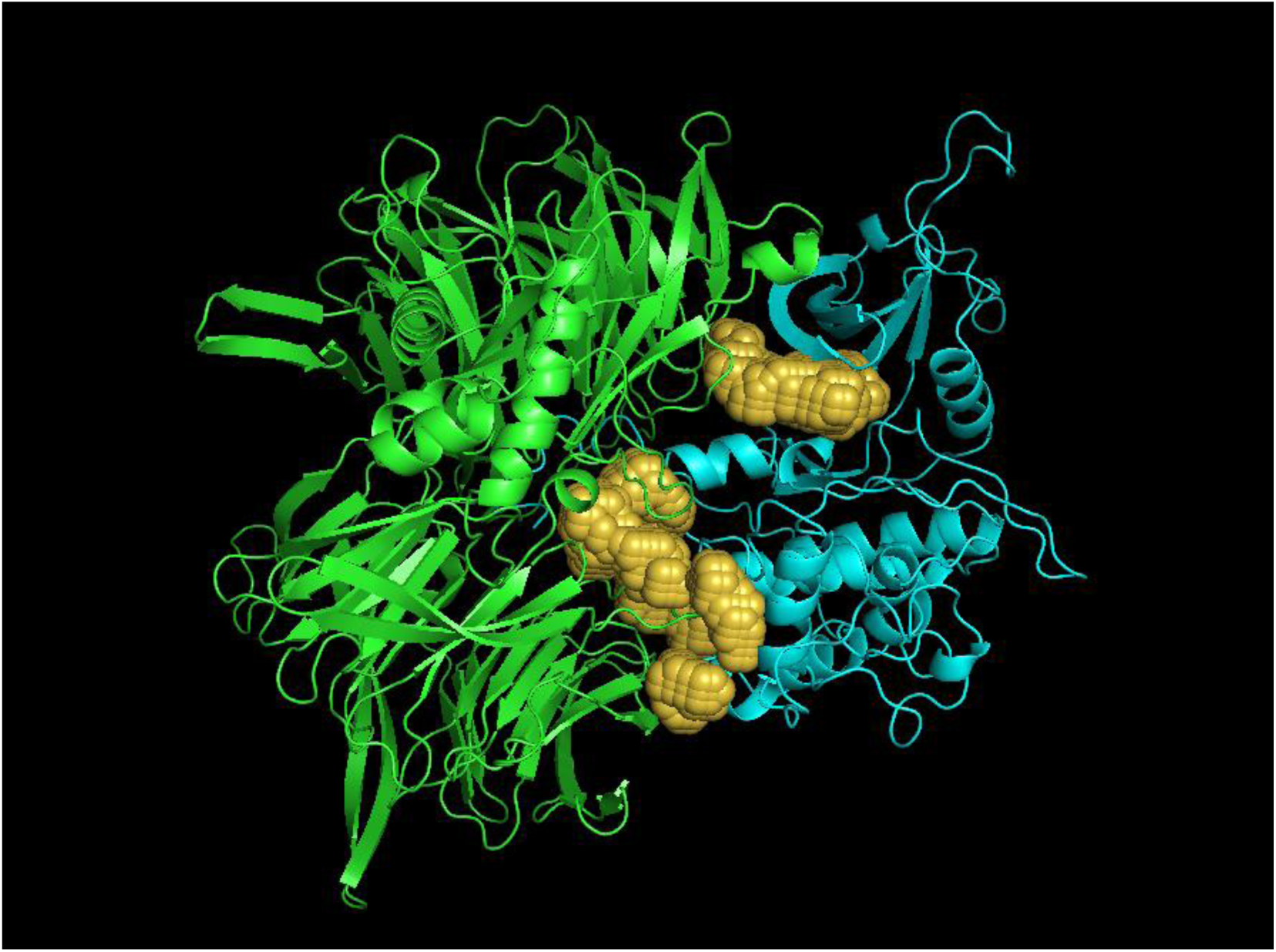
Analysis of the pocket formation at the interface of the two proteins for PPI_Pose_2.

**Table 1.** Protein-Protein docking and interface pocket compatibility for molecular glue binding.

| Pose ID | Protein-Protein Interaction (PPI) score from Megadock | Interface pocket compatibility |
| --- | --- | --- |
| PPI_Pose_1 | 4441 | No |
| PPI_Pose_2 | 4001 | Yes |
| PPI_Pose_3 | 3883 | No |
| PPI_Pose_4 | 3779 | No |
| PPI_Pose_5 | 3523 | No |
| PPI_Pose_6 | 3479 | No |
| PPI_Pose_7 | 3421 | No |
| PPI_Pose_8 | 3397 | No |
| PPI_Pose_9 | 3319 | No |
| PPI_Pose_10 | 3307 | No |

Further, at the pocket formed at the interface of these two proteins, molecular docking was carried out using AutoDock-Vina which reproduced the experimental pose of the molecular glue (RC8) in PDB ID: 6TD3 as shown below with docking derived pose in yellow and experimental pose in cyan in Figure 4.

**Figure 4.**
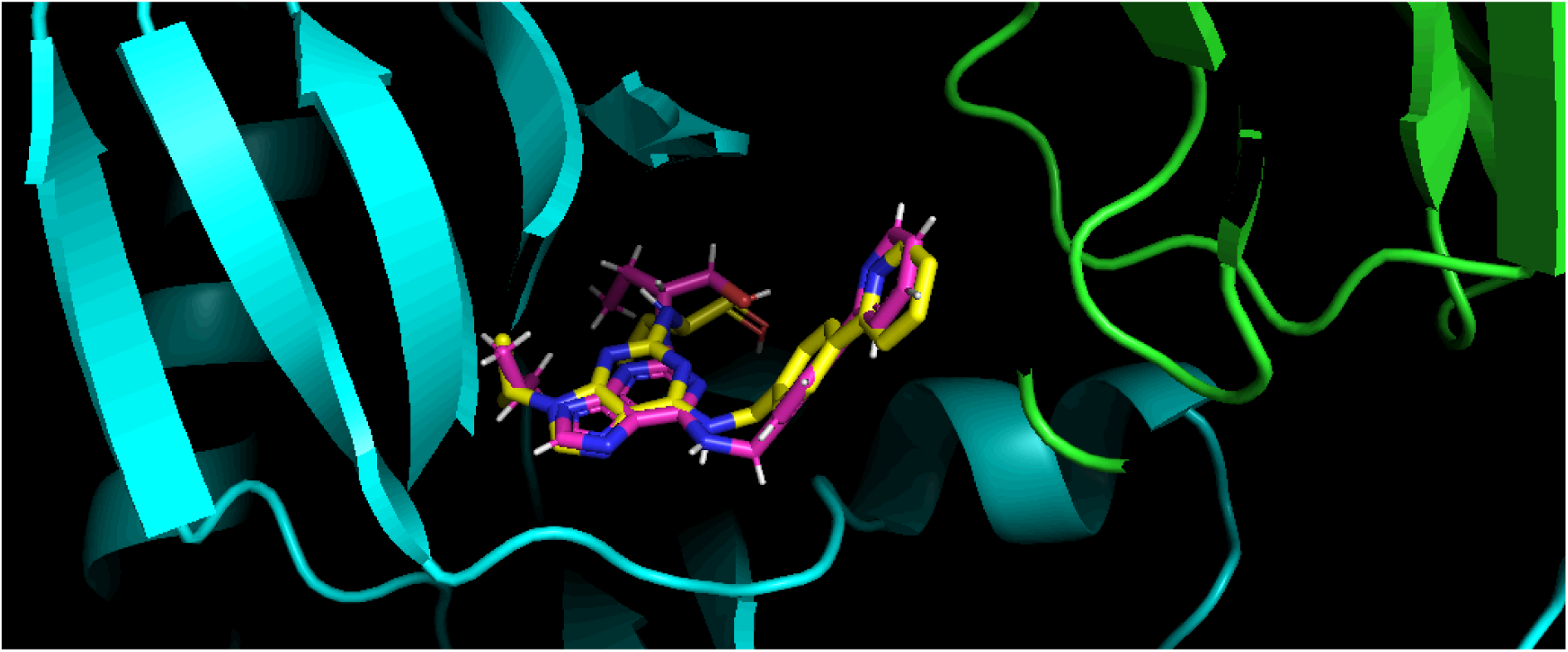
Pose of RC8 reproduced in DDB1-CDK12 and the resulting ternary complex.

### 2. Molecular glue mediated favorability of the two proteins to form a ternary complex

The diversity of interactions mediated by molecular glues in ternary complexes reported in RCSB PDBs in the list collected by Haun et. al [12] was studied using LigPlot+ [13] and it was found that molecular glues can mediate either covalent or non-covalent interactions and a representative example for each case is discussed below along with the interactions from LigPlot. For the covalent case, PDB ID: 8G46 was chosen as the representative example for the class and for non-covalent case PDB ID: 6TD3 was chosen as the representative example for the class.

As seen below in Figure 5, covalent interactions are manifest in PDB ID: 8G46 system, wherein the Cysteine residue (Cys58) a usual participant of covalent bonding interaction is observed to interact with the molecular glue (YK3).

**Figure 5.**
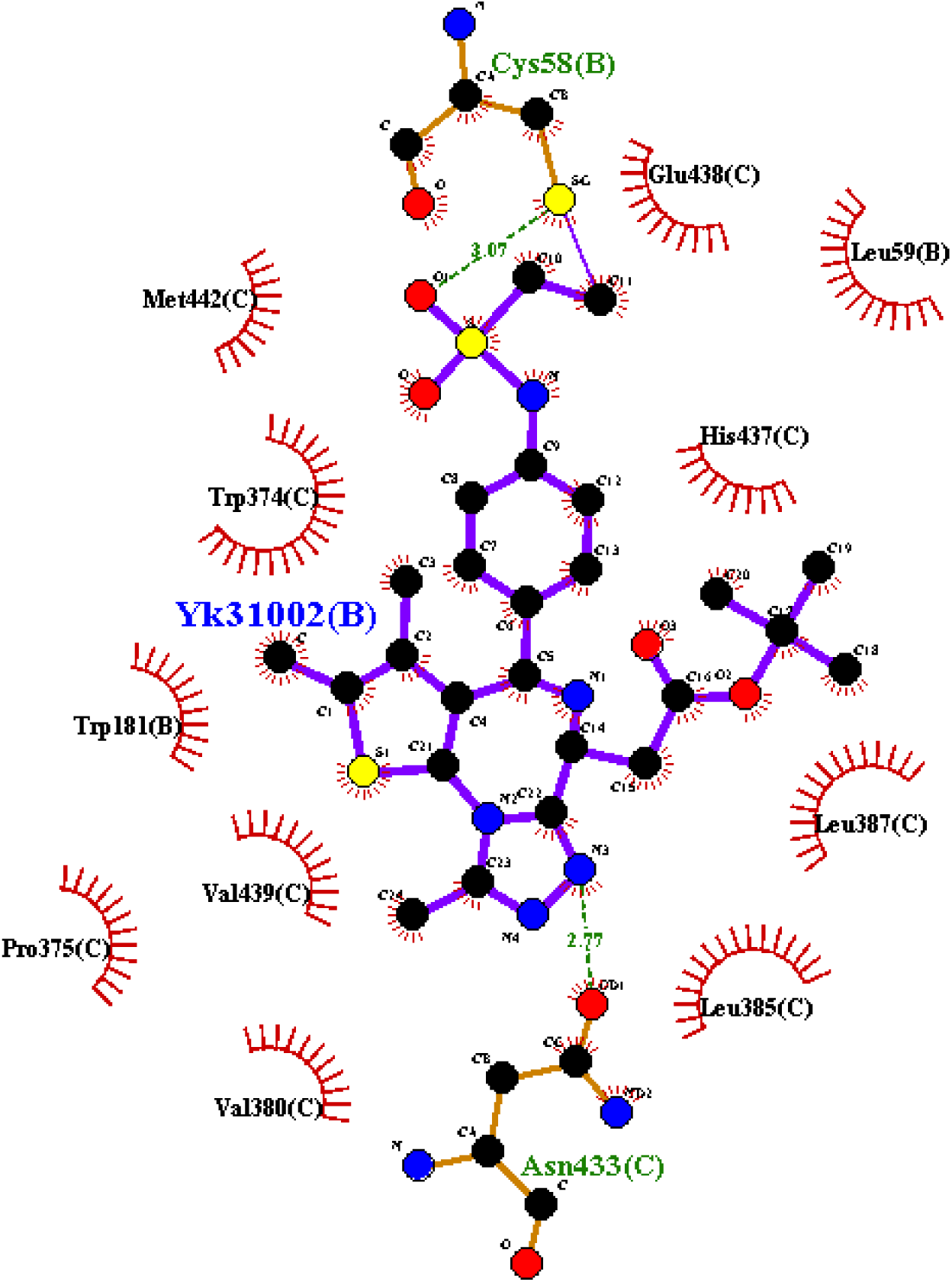
YK3 as representative example for molecular glues that work through covalent interactions.

Only non-covalent interactions are observed in the PDB ID: 6TD3 system where molecular glue (RC8) has hydrophobic interactions with the hydrophobic residues in the pocket and hydrogen bonding with MET816 and ASP819 as shown in Figure 6 below.

**Figure 6.**
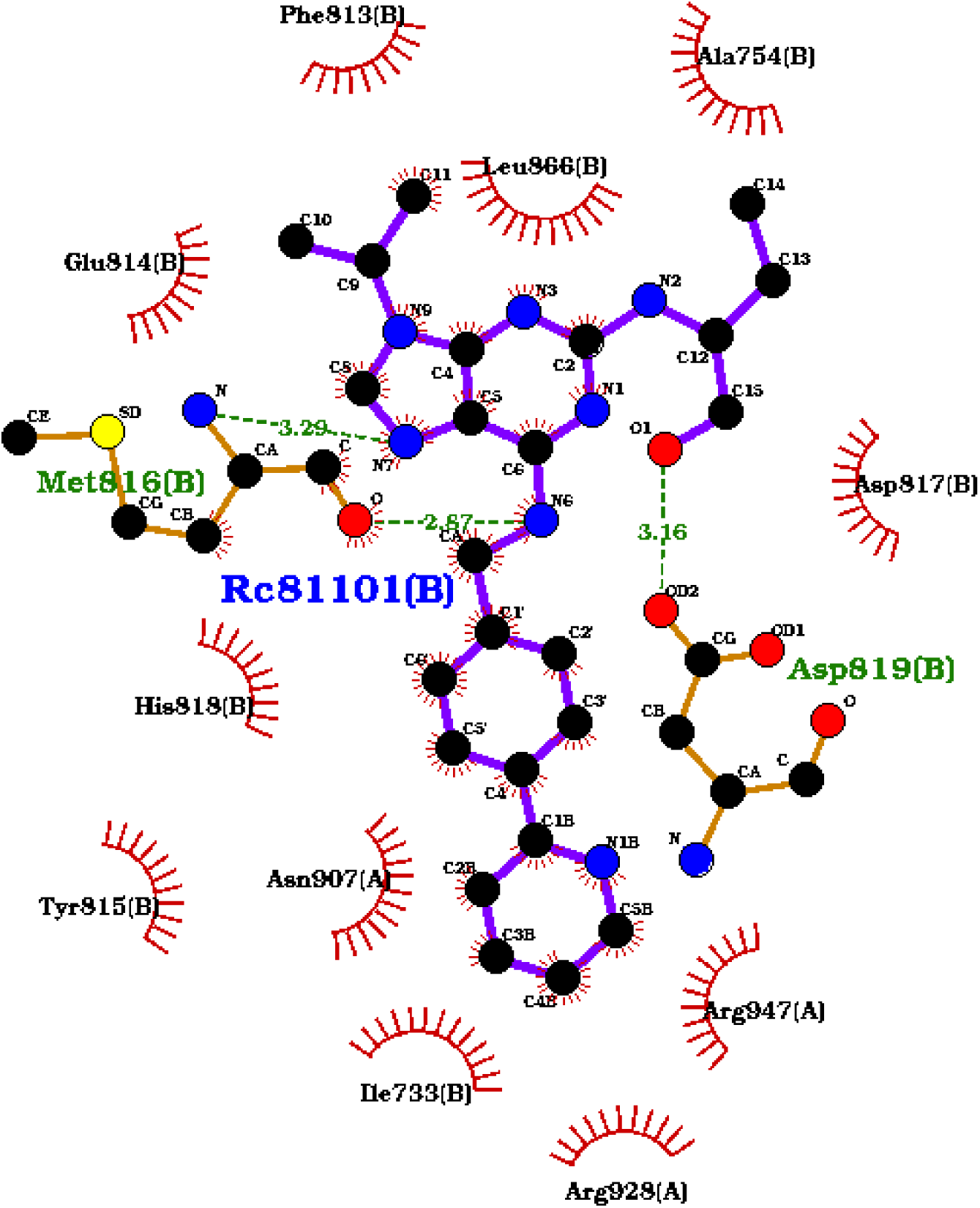
RC8 as representative example for molecular glues that work through non-covalent interaction.

The molecular glue induced thermodynamic favourability to induce the Protein-Protein interaction in the systems associated with PDB IDs 8G46, 7TE8 and 6TD3 was assessed using theoretical ΔG calculations carried out using MMPBSA method. A classical molecular dynamics simulation was carried out for 50 ns for the protein-protein system associated with PDB IDs 8G46, 7TE8 and 6TD3 with and without the molecular glue and the stabilization of the protein-protein complex was observed using the RMSD stabilization graphs shown in Figures 7, 8, and 9 below and the stable portion of the trajectory was used for MMPBSA based ΔG calculations. Results of the MMPBSA calculations are tabulated in Table 2.

**Figure 7.**
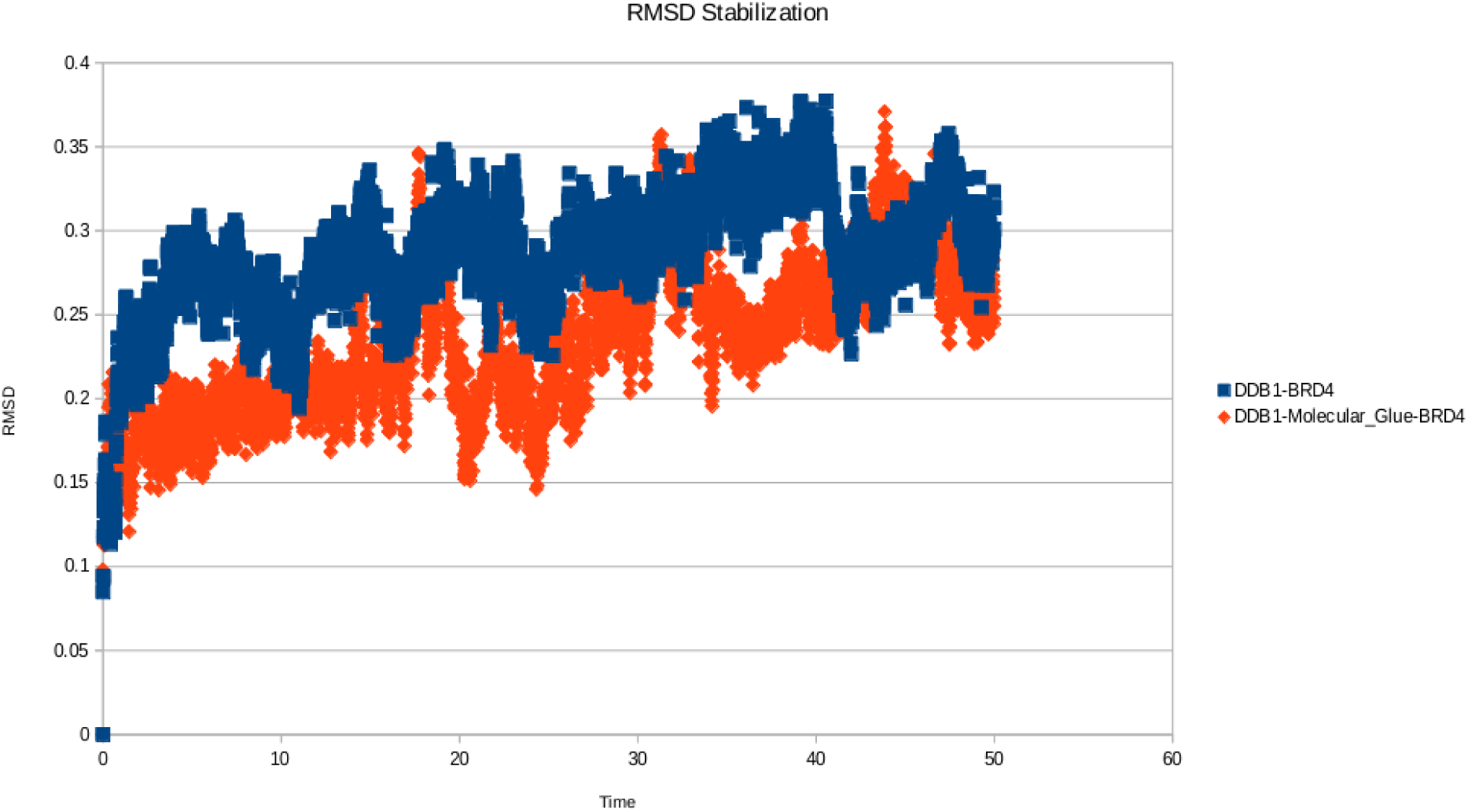
RMSD Stabilization of PDB 8G46 system with and without the molecular glue.

**Figure 8.**
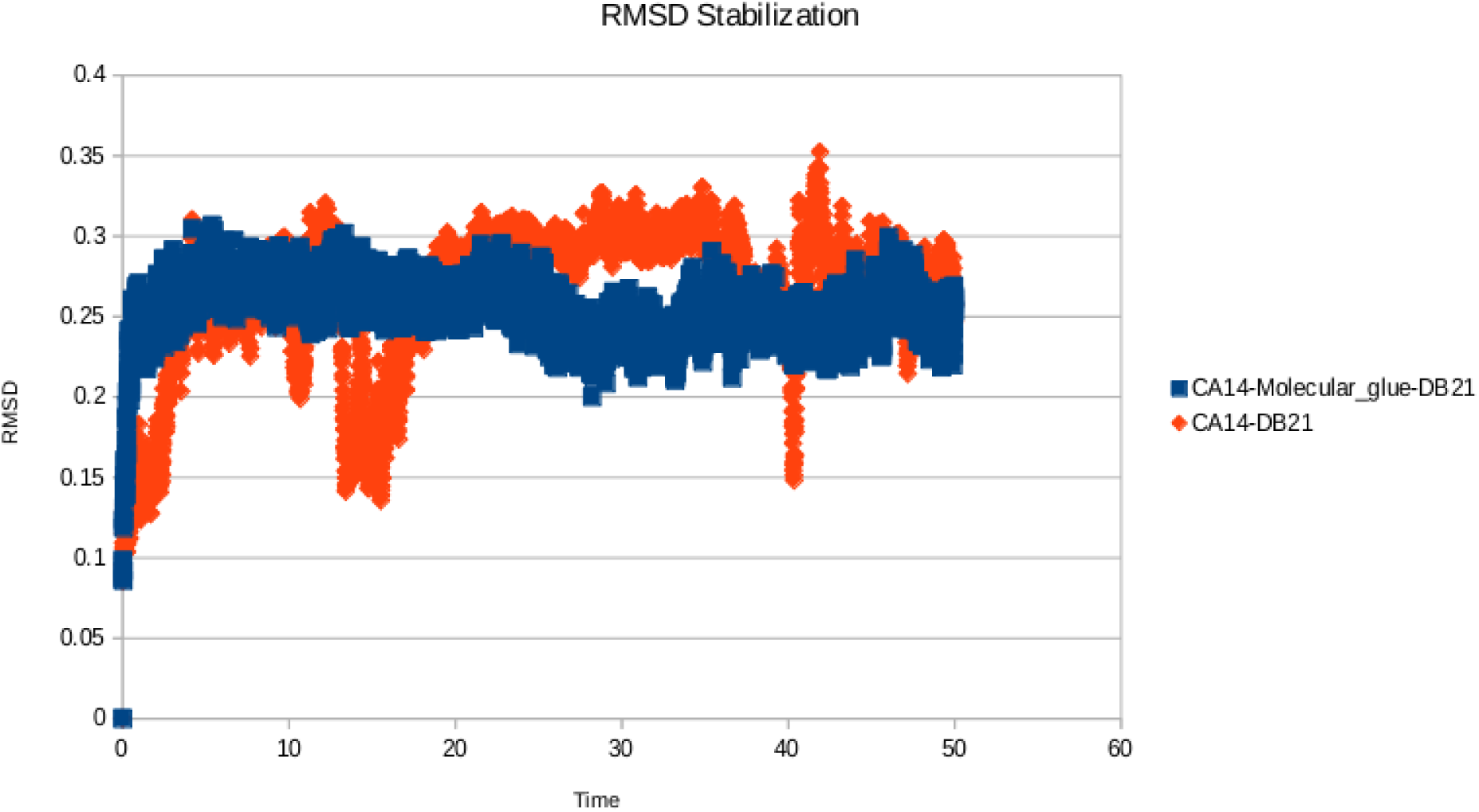
RMSD Stabilization of PDB 7TE8 system with and without the molecular glue.

**Table 2.** MMPBSA based calculations.

| System | Complex | MMPBSA based $\Delta G$ (kcal/mol) | Receptor (R) and Ligand (L) definitions for MMPBSA calculations |
| --- | --- | --- | --- |
| PDB ID: 8G46 | DDB1-BRD4 | -46.3 | R=BRD4<br>L=DDB1 |
|  | DDB1-Molecular_glue-BRD4 | -64.7 | R=BRD4<br>L=DDB1-Molecular_glue |
| PDB ID: 7TE8 | CA14-DB21 | -27.8 | R=CA14<br>L= DB21<br>L=DB21-Molecular_glue |
|  | CA14-Molecular_glue-DB21 | -119.1 | R=CA14<br>L=DB21-Molecular_glue |
| PDB ID: 6TD3 | DDB1-Cyclin_dependent_kinase_12 | -47.1 | R=DDB1<br>L=Cyclin_dependent_kinase_12 |
|  | DDB1-Molecular_Glue-Cyclin_dependent_kinase_12 | -73.19 | R=DDB1 |
|  |  |  | L=<br>Cyclin_dependent_kinase_12-Molecular_Glue |

From the results of the MMPBSA calculations shown in Table 2, it can be inferred that the molecular glue enhanced the protein-protein interaction in all 3 cases.

Furthermore, molecular glues are also known to enhance protein-protein interaction through molecular glue mediated conformation change as in Thalidomide mediated conformation change of CRBN that enhances interaction between CRBN and Casein Kinase 1α (CK1α) interaction leading to subsequent degradation of CK1α [22, 23]. To investigate this mechanism in-silico, an ensemble protein-protein interaction poses was generated using AlphaFlow and, the molecular glue (Thalidomide) stabilizing the closed (Holo) form of CRBN as opposed to its open (Apo) form which results in increase in interaction between CRBN and CK1α was rationalized through the scores obtained which are tabulated in Table 3 below. The generated ensembles of conformation were compared against the experimentally known Open (Apo) form of CRBN as in PDB ID: 8D7X and Closed (Holo) form as in PDB ID: 5FQD.

**Table 3.** In-silico investigation of CRBN-CK1α allostery mediated by molecular glue (Thalidomide)

| PPI poses of CRBN-CK1α | RMSD from Alignment of PPI pose with CRBN Open (Apo) form PDB | RMSD from Alignment of PPI pose with CRBN Closed (Holo) form | PPI interaction of CRBN-CK1α from SURFACE | Protein-ligand interaction score of Molecular Glue (Thalidomide) and CRBN |
| --- | --- | --- | --- | --- |
| **Pose1** | **3.570** | **0.874** | **-33.89** | **-7.89** |
| Pose2 | 2.234 | 1.113 | -23.89 | -6.15 |
| Pose3 | 2.781 | 1.783 | -25.41 | -6.31 |
| Pose4 | 1.435 | 2.187 | -25.78 | -6.81 |
| Pose5 | 1.133 | 2.257 | -26.11 | -6.07 |
| **Pose6** | **0.759** | **4.112** | **-21.17** | **-5.77** |
| Pose7 | 1.738 | 3.071 | -17.88 | -5.89 |
| Pose8 | 2.178 | 4.311 | -16.81 | -6.07 |
| Pose9 | 3.119 | 2.788 | -20.11 | -5.68 |
| Pose10 | 4.148 | 3.791 | -18.81 | -5.57 |

It can be inferred from above Table 3 that the molecular glue Thalidomide binds to CRBN and stabilizes the closed form of CRBN and the closed form of CRBN has an enhanced interaction with CK1α as compared to the open form. The MMPBSA scores shown below in Table 4 further support the same inference.

**Table 4.** In-silico investigation of CRBN-CK1α allostery mediated by molecular glue (Thalidomide) through MMPBSA based calculations.

| System | | $\Delta G$ (kcal/mol) |
| --- | --- | --- |
| CRBN- CK1α | CRBN closed form | -67.89 |
|  | CRBN open form | -44.13 |

The open and closed form of CRBN bound to CK1α is shown in Figure 10 below with the open form in green and the closed form (in purple) stabilized by the molecular glue (in cyan) having an enhanced interaction with CK1α (in yellow)

**Figure 9.**
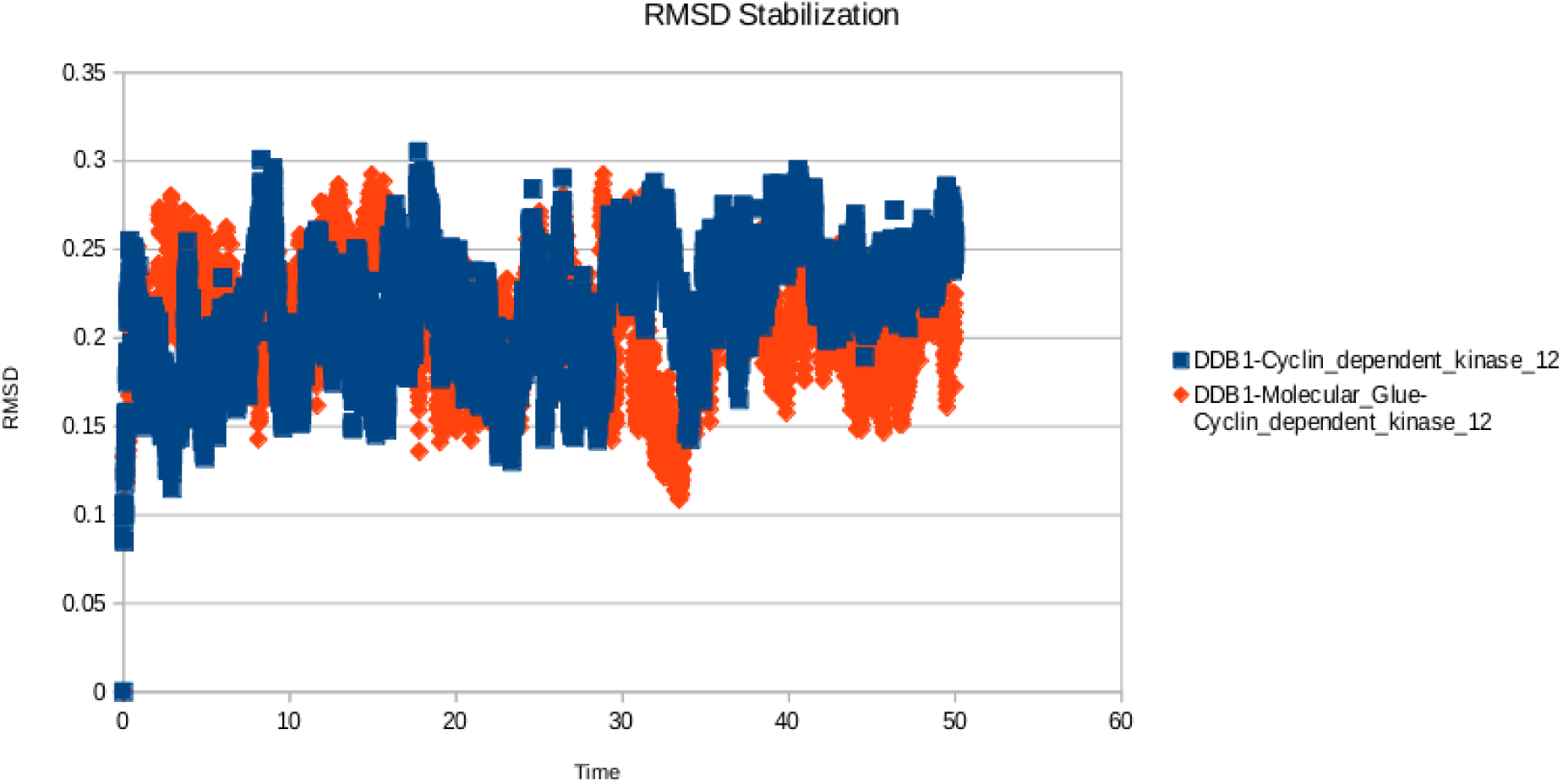
RMSD Stabilization of PDB 6TD3 system with and without the molecular glue.

**Figure 10.**
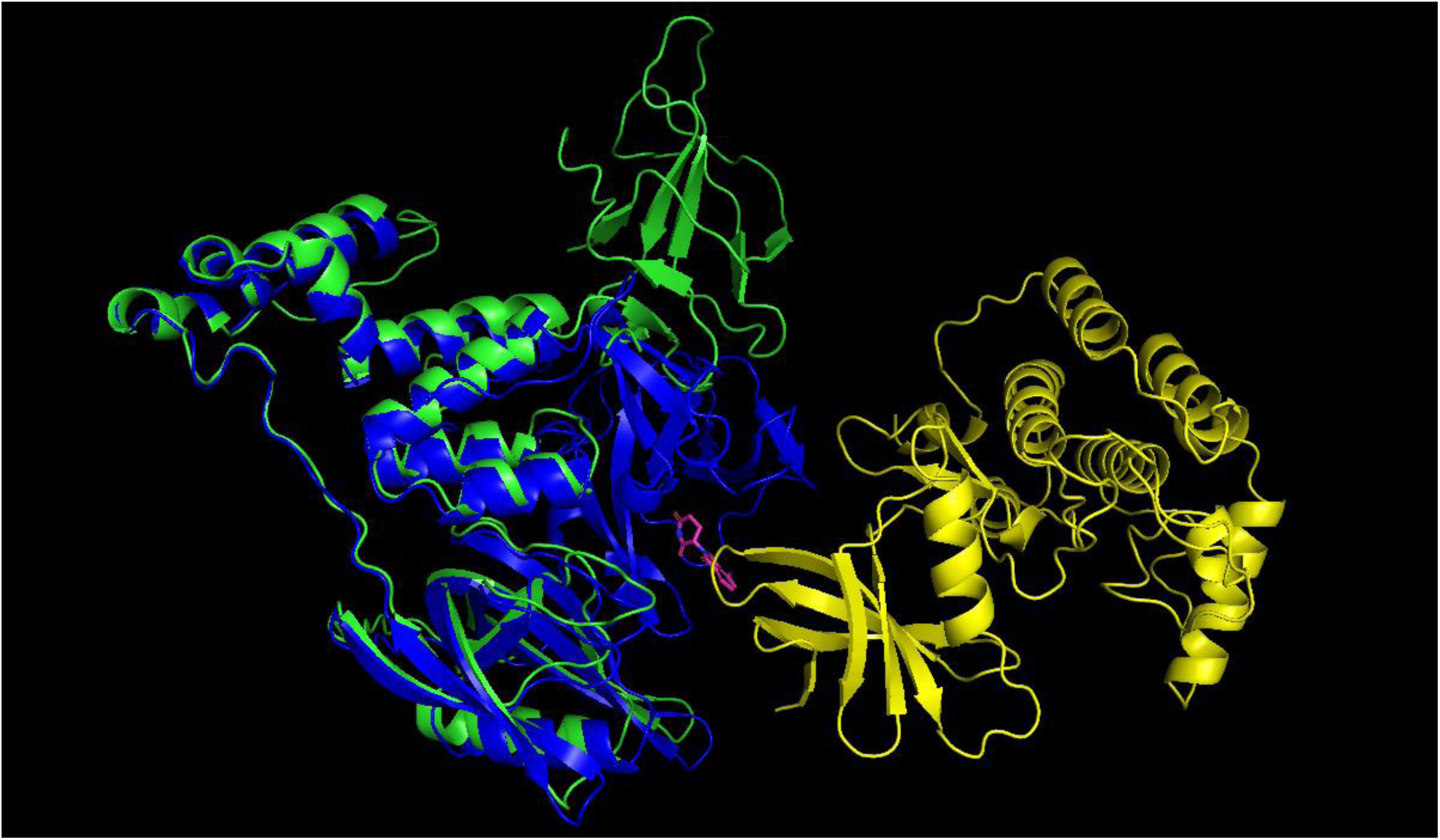
Molecular glue allostery mediated CRBN-CK1α interaction.

**Figure 11.**
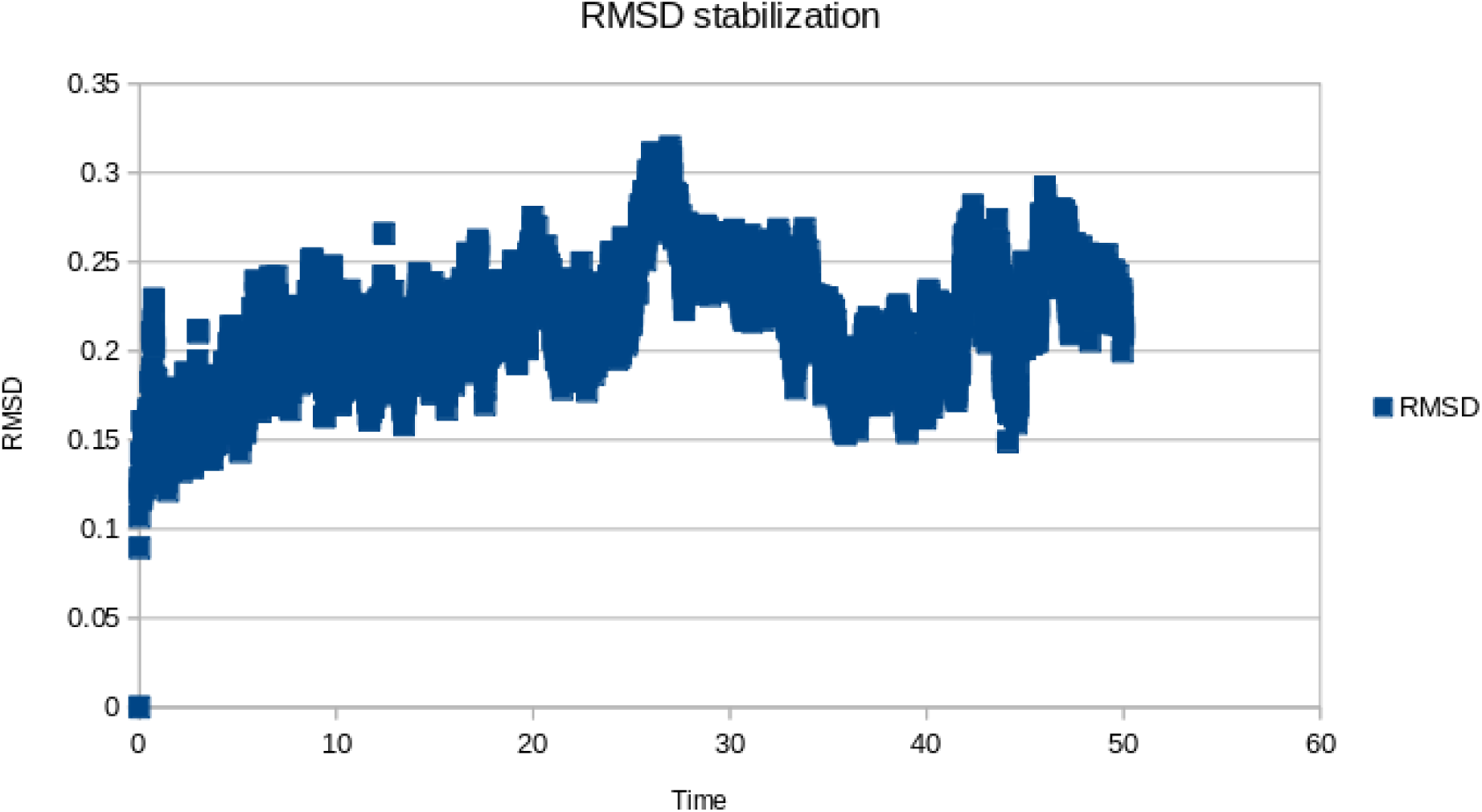
RMSD stabilization of DDB1-Cyclin_dependent_kinase_12 ternary complex stabilized by SAIT_MG_26121.

### 3. In-silico differentiation of the stability and ternary binding constants (K_d_) associated with the molecular glue mediated ternary complex

Free energy perturbation and QM/ MM calculations were carried out on two systems with PDB ID: 6H0G and 7BQV which have a micromolar ternary binding constants (K_d_) of 2.3 µM for 6H0G and a nanomolar ternary binding constants (K_d_) of 1.8 nM for 7BQV. Free energy perturbation calculations and QM/ MM calculations show that it is possible to differentiate the ternary systems of different orders of ternary binding constant values as shown in Table 5.

**Table 5.**
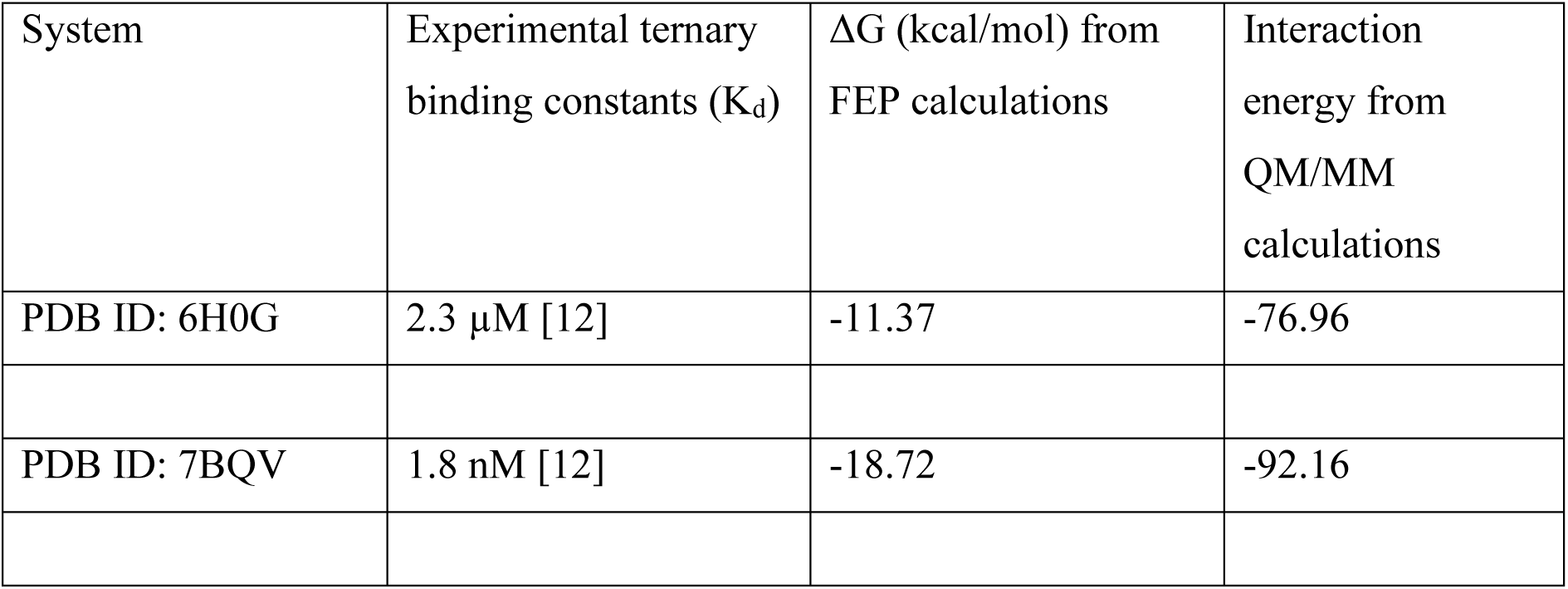
FEP and QM/ MM calculations to differentiate stability of molecular glue mediated ternary complex.

Further, Thermal Titration Molecular Dynamics (TTMD) was also carried out to estimate retention of interaction in an increasing temperature ramp. Results obtained are presented below in Table 6.

**Table 6.** TTMD calculations to differentiate stability of molecular glue mediated ternary complex.

|  | Temperature Ramp |  |  |  |  |
| --- | --- | --- | --- | --- | --- |
|  | 300K |  | 450K |  | Remarks and Interpretation |
|  | **K<sub>D</sub>(Ternary)=2.3 μM**<br>PDB ID: 6H0G | **K<sub>D</sub>(Ternary)=0.0018 μM**<br>PDB ID: 7BQV | **K<sub>D</sub>(Ternary)=2.3 μM**<br>PDB ID: 6H0G | **K<sub>D</sub>(Ternary)=0.0018 μM**<br>PDB ID: 7BQV |  |
| Average interaction retained | 0.71 | 0.83 | 0.07 | 0.29 | Should be higher for tight binder than weak binder. |

The obtained TTMD profile indicates that the ternary system mediated by the molecular glue in 7BQV is more stable as compared to 6H0G which is consistent with the experimentally known K_D_ (Ternary) for 7BQV and 6H0G, respectively.

### 4. AI driven de novo design of molecular glues for PDB ID: 6TD3 system

AI driven de novo molecular design approaches [21] were employed to design new molecular glues for the PDB ID: 6TD3 system around the chemical space of the base scaffold derived from RC8. RC8 is a potent nano molar compound, so we aimed to design a molecular glue which will be comparable to RC8, if not better. The newly designed molecular glues were screened through the developed approach and results are tabulated below. The newly designed molecular glues were docked in the pocket of RC8 and the interactions of RC8 and the new designs are tabulated in Table 7. The interactions conserved among RC8, and the new designs are highlighted in green and based on the SURFACE protein-ligand interaction score, the candidate ‘SAIT_MG_26121’ was shortlisted for further consideration.

**Table 7.**
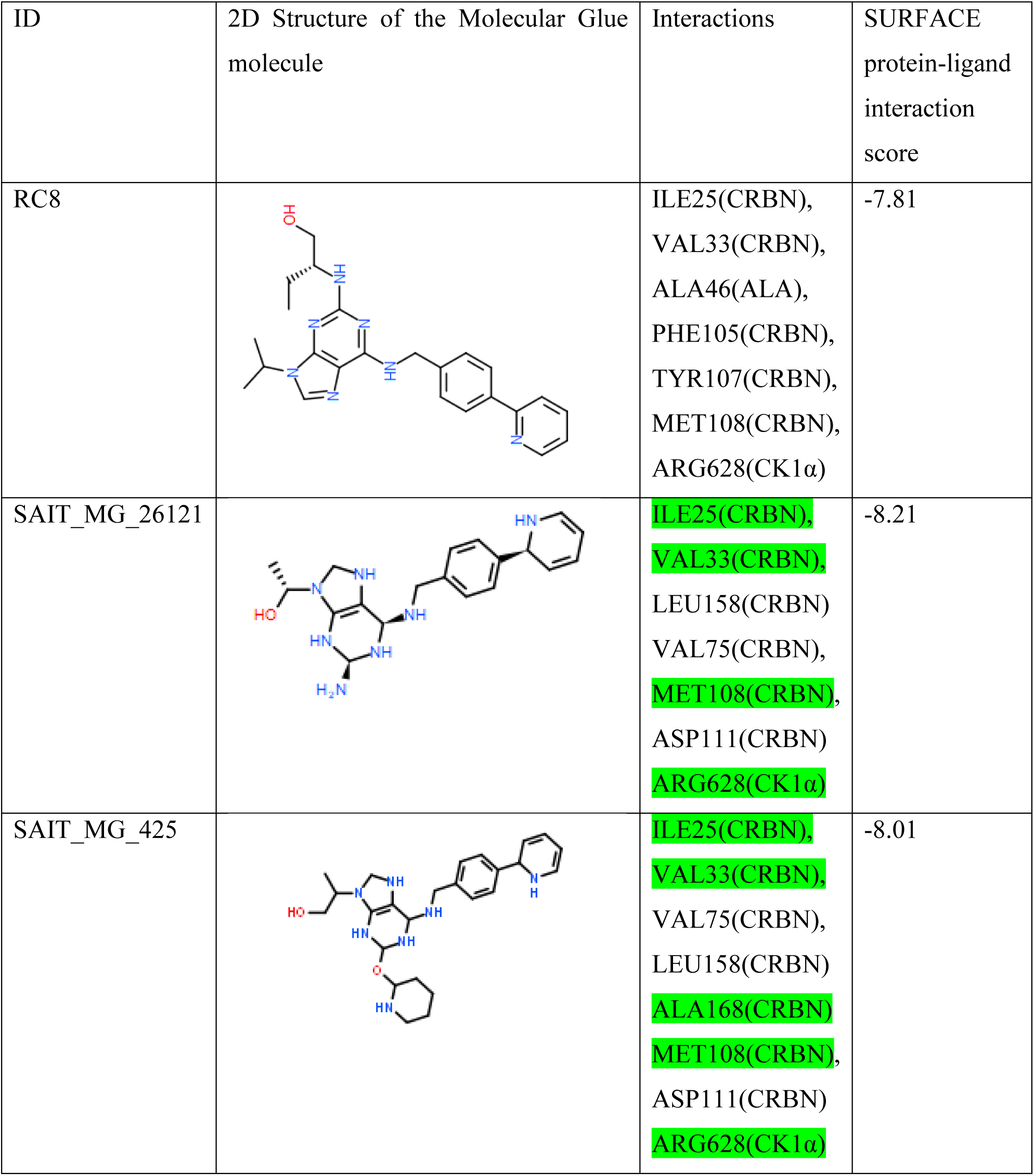
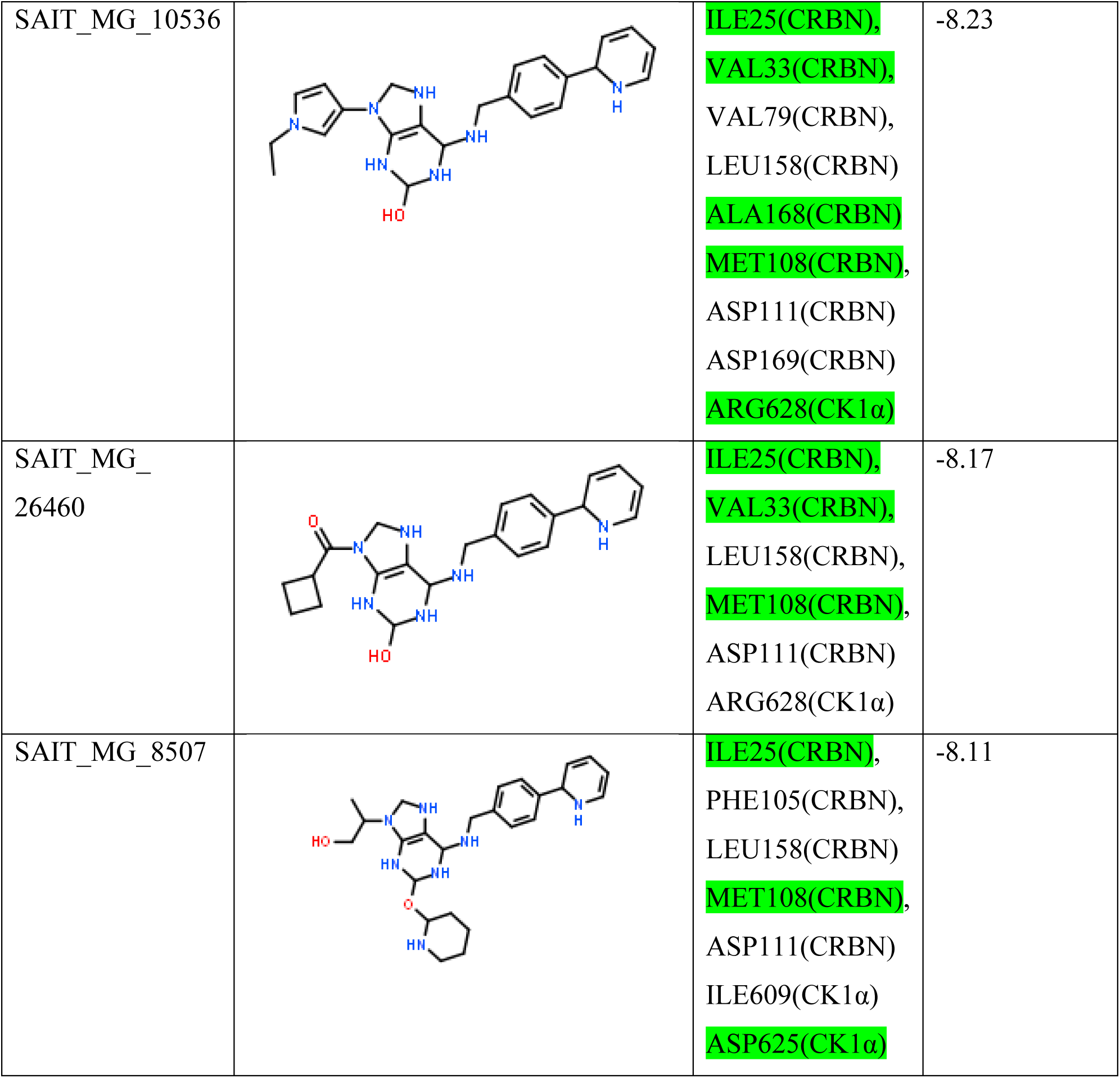
De novo design of molecular glues for 6TD3 system.

| ID | 2D Structure of the Molecular Glue molecule | Interactions | SURFACE protein-ligand interaction score |
| --- | --- | --- | --- |
| RC8 | | ILE25(CRBN),<br>VAL33(CRBN),<br>ALA46(ALA),<br>PHE105(CRBN),<br>TYR107(CRBN),<br>MET108(CRBN),<br>ARG628(CK1α) | -7.81 |
| SAIT_MG_26121 | | ILE25(CRBN),<br>VAL33(CRBN),<br>LEU158(CRBN)<br>VAL75(CRBN),<br>MET108(CRBN),<br>ASP111(CRBN)<br>ARG628(CK1α) | -8.21 |
| SAIT_MG_425 | | ILE25(CRBN),<br>VAL33(CRBN),<br>VAL75(CRBN),<br>LEU158(CRBN)<br>ALA168(CRBN)<br>MET108(CRBN),<br>ASP111(CRBN)<br>ARG628(CK1α) | -8.01 |
| SAIT_MG_10536 | | ILE25(CRBN),<br>VAL33(CRBN),<br>VAL79(CRBN),<br>LEU158(CRBN)<br>ALA168(CRBN)<br>MET108(CRBN),<br>ASP111(CRBN)<br>ASP169(CRBN)<br>ARG628(CK1α) | -8.23 |
| SAIT_MG_26460 | | ILE25(CRBN),<br>VAL33(CRBN),<br>LEU158(CRBN),<br>MET108(CRBN),<br>ASP111(CRBN)<br>ARG628(CK1α) | -8.17 |
| SAIT_MG_8507 | | ILE25(CRBN),<br>PHE105(CRBN),<br>LEU158(CRBN)<br>MET108(CRBN),<br>ASP111(CRBN)<br>ILE609(CK1α)<br>ASP625(CK1α) | -8.11 |

Further, MMPBSA calculations based on binding free energy calculations were carried out to understand the molecular glue (SAIT_MG_26121) induced favourability for ternary complex formation for the DDB1-Cyclin_dependent_kinase_12 system. A 50 ns classical molecular dynamics simulation was carried out to estimate the stability of the ternary complex mediated by SAIT_MG_26121. RMSD stabilization is shown in Figure11 below. The stable portion of the trajectory was chosen for MMPBSA based binding energy calculations which indicate the molecular glue (SAIT_MG_26121) induced ternary complexation and the results are tabulated in Table 8 below.

**Table 8.** MMPBSA calculations for ‘SAIT_MD_26121’.

| System | | MMPBSA $\Delta G$ (kcal/mol) |
| --- | --- | --- |
| PDB ID: 6TD3 | DDB1-Cyclin_dependent_kinase_12 | -47.1 |
| SAIT_MD_26121 | DDB1-Molecular_Glue-Cyclin_dependent_kinase_12 | -74.21 |

Finally, the stability of the molecular glue mediated ternary complex mediated by RC8 (reference) and the de novo designed candidate ‘SAIT_MD_26121’ was compared through FEP and QM/ MM scores which are tabulated in Table 9 and their respective TTMD profiles which are tabulated in Table 10.

**Table 9.** FEP and QM/ MM calculations for SAIT_MD_262121.

| System | Experimental ternary binding constants ($K_d$) | $\Delta G$ (kcal/mol) from FEP calculations | Interaction energy from QM/MM calculations |
| --- | --- | --- | --- |
| PDB ID: 6TD3 (RC8) | < 100 nM [12] | -17.11 | -105.22 |
| SAIT_MD_21621 |  | -17.89 | -103.81 |

**Table 10.** TTMD profile for SAIT_MD_26121.

|  | Temperature Ramp |  |  |  |  |
| --- | --- | --- | --- | --- | --- |
|  | 300K |  | 450K |  | Remarks and Interpretation |
|  | RC8 | SAIT_MD_26121 | RC8 | SAIT_MD_26121 |  |
| Average interaction retained | 0.73 | 0.69 | 0.17 | 0.19 | SAIT_MD_26121 retains interactions similar to reference RC8 at higher temperature indicating a strong binding profile. |

The results of the FEP, QM/ MM and TTMD calculations indicate that newly designed candidate ‘SAIT_MD_26121’ has a similar binding profile as that of the reference RC8 which is itself a nano molar molecular glue.

## Conclusions

Molecular glues, which more closely resemble traditional small molecules, offer a promising alternative to heterobifunctional PROTACs for target protein degradation. The PROTAC field faces challenges related to permeability and bioavailability due to the large size of PROTAC molecules. However, for serendipitously discovered molecular glues, a rational design approach remains elusive.

To establish the in-silico rationale for molecular glue design, we take known molecular glue mediated ternary complexes reported in RCSB PDB and develop in-silico methods that are able to do the following 1) Reproduce experimentally known binding modes of molecular glues and the ternary complex formed therein, 2) Rationalize the thermodynamic favorability induced by molecular glues for ternary complex through theoretical ΔG calculations, 3) Differentiate stability of molecular glue mediated ternary complexes. After carrying out a retrospective validation for the developed approach, we use the developed approach to design a new molecular glue as a demonstrative case. By bridging the gap in rational in-silico design for molecular glues, we are aiming here to contribute valuable insights to the literature and provide a foundation for future development and improvements.

## Acknowledgements

We are grateful for the various useful discussions we have had with our colleagues at Sravathi AI Technology Pvt Ltd. In particular, we would like to thank Raghu Bhagavat, Srinivasan Krishnaswami, Nivedita Bharti, and Rajesh Kondabala for their valuable input and insights.

## Data availability and reproducibility statement

All results can be reproduced as per the methodology reported in our methodology section and the use of publicly available research data and software packages which are as follows: Data:

[1] All molecular glue ternary complexes used as a part of developing the methodology were download from RCSB PDB – https://www.rcsb.org/.

Software packages:

[2] MEGADOCK for protein-protein docking (https://github.com/akiyamalab/MEGADOCK).

[3] CAVIAR for pocket identification CAVIAR for Pocket identification (https://github.com/jr-marchand/caviar).

[4] RDKIT for cheminformatics tasks (https://github.com/rdkit/rdkit).

[5] For molecular dynamics simulations involving Ternary complex modelling, TTMD, and Free energy calculations, we used GROMACS 2023 software (https://github.com/gromacs/gromacs).

[6] For MMPBSA based free energy calculations, we used GMX_MMPBSA software (https://github.com/Valdes-Tresanco-MS/gmx_MMPBSA).

[7] For Interaction Fingerprinting in TTMD, we used PLIP (https://github.com/pharmai/plip).

[8] ORCA for QM calculations (https://www.faccts.de/orca/).

